# (1→3)-β-D-glucan derivatives with unique structural properties differentially affect murine lung inflammation and histology

**DOI:** 10.1101/2023.10.18.562993

**Authors:** N. Metwali, E.M. Stapleton, S. Hadina, P.S. Thorne

## Abstract

**Background:** β-D-glucan (BDG) compounds are proinflammatory fungal cell wall polysaccharides known to cause respiratory pathology. However, the specific immunological effect of unique BDG structures on pulmonary inflammation is understudied. We sought to characterize the effect of four unique fungal BDGs with unique branching patterns, solubility, molecular weights and higher order structure in murine airways.

**Methods:** Scleroglucan (1→3) (1→6)-highly branched BDG, laminarin (1→3) (1→6)-branched BDG, curdlan (1→3)-linear BDG, and pustulan (1→6)-linear BDG were assessed by nuclear magnetic resonance spectroscopy (NMR). Glucan compounds were confirmed negative for endotoxin by kinetic chromogenic Limulus amebocyte lysate assay. Each BDG was tested using an inhalation model with C3HeB/FeJ mice and compared to control mice exposed to saline and unexposed sentinels (all, n=3-19). Inhalation studies were performed using all BDGs ±heat-inactivation (1-hr. autoclave) given BDG solubility increases after heat-inactivation. Outcomes included bronchoalveolar lavage (BAL), differential cell counts (macrophages, neutrophils, lymphocytes, and eosinophils), BALF cytokine and serum IgE and IgG2a expression (by multiplex immunoassay and ELISA). Lastly, *ex vivo* cultured left lung cells were BDG stimulated, and cytokines (multiplex) compared to unstimulated cells. Right lung histology was performed.

**Results:** While each BDG affected lung inflammatory outcomes, pustulan exposure led to the most pro-inflammatory profile of all BDGs tested, including increased inflammatory infiltrate into BAL, increased serum IgE and IgG2a and increased cytokine production. Secondary stimulation of sensitized lung cells by lipopolysaccharide (LPS) significantly increased cytokine expression. Lung histology revealed significant fibrosis in lungs exposed to soluble BDGs pustulan and scleroglucan.

**Conclusion:** Inhalation of BDGs with distinct branching patterns exhibited varying inflammatory potency and immunogenicity, which lichen-derived pustulan having the greatest effect. Glucan source and solubility should be considered in exposure and toxicological studies.

## INTRODUCTION

Damp and moldy environments are known contributors to respiratory pathology, including allergic asthma, in children and adults (1–4). In children, for example, increased exposure to fungal cell wall material, β-D-glucan (BDG), is negatively associated with FEV_1_ and increases odds of emergency/urgent care visits in asthmatics by nearly nine-fold (5). Exposure to indoor bacterial endotoxin and mold BDG are known respiratory irritants and potent cytokine inducers (6–8).

β-D-glucan structures vary depending on their source and extraction process (9, 10). (1→3)-β-D-glucans (1,3-BDGs) are major structural components of fungal cell walls that activate macrophages and modulate innate immunity. Microorganism-derived BDGs generally have (1→ 3)-linked anhydro-D-glucose units as a backbone with periodic (1→ 6)-linked side chains. Other glucans have linear (1→ 3) linkage without branching. Numerous studies have revealed critical BDG properties include their molecular weight (MW) and water solubility, which depend on their degree of branching (DB) (11–13). Glucans are typically most potent between 100-200 kDa with the DB ranging from 0.2-0.5 (10). Various destructive and nondestructive methods are available to define BDG molecular structure, DB, and degree of polymerization. Non-destructive light-scattering techniques are sensitive to macromolecular aggregation and provide glucan NMR and polydispersity (11).

Glucan properties have downstream immunomodulatory effects. For example, BDG receptor Dectin-1, a non-toll-like receptor (TLR) pattern-recognition receptor (PRR), uniquely recognizes soluble and particulate glucans; Dectin-1 signaling is antagonized by soluble glucans (14, 15). BDG properties can be exploited therapeutically as biological-response modifiers to improve host immunity (16–18). For example, branched 1,3-BDGs including lentinan (19), schizophyllan (20), and krestin (21) provide antitumor activity in the host. Soluble glucans are also thought to have unique anti-tumor effects (22). Yeast cell wall-derived zymosan is a potent macrophage activator and neutrophil chemoattractant, binding to membrane components such as scavenger receptor complement receptor 3, Lactosylceramide, and Dectin-1 with TLR2 and TLR6 heterodimer signaling (23–26). Linear 1,3-BDGs, (e.g., curdlan) can also stimulate macrophage binding to pattern-recognition receptors using MyD88 for signal transduction (28).

Specific 1,3-BDG physicochemical parameters, including primary structure, solution conformation, MW and/or polymer charge are thought to affect macrophage receptor affinity. However, this relationship has not been defined, in part due to the lack of well-characterized 1,3-BDG polymers with varying MWs and conformations. Their effect on respiratory inflammation is understudied. Therefore, we evaluated the effect of four glucan compounds on lung inflammation using an *in vivo* murine allergen inhalation model.

## MATERIALS AND METHODS

### Glucan compounds

Scleroglucan (1→3) (1→6)-highly branched BDG was generously provided by Dr. David Williams (Tennessee State University). Commercial suppliers provided laminarin ((1→3) (1→6)-branched BDG; Sigma-Aldrich, Inc., St. Louis, MO), curdlan ((1→3)-linear BDG; Wako, Richmond, VA), and pustulan ((1→6)-linear BDG; Calbiochem Inc.,La Jolla, CA). The degree of branching was determined by nuclear magnetic resonance (NMR), **Appendix Methods**.

### Animal model

Four-to five-week-old, pathogen-free male C3HeB/FeJ mice were obtained from Jackson Laboratories (Bar Harbor, ME) and housed in a rodent vivarium with a 12h light-dark cycle and provided with food and water *ad libitum*. Mice were quarantined for 10 days prior to BDG glucan exposure. All animal protocols were reviewed and approved by the Office of the Institutional Animal Care and Use Committee at the University of Iowa.

### β-D-glucan conjugation

We performed BDG conjugation using previously described methods (29–32). Curdlan and pustulan were oxidized using 2 mL of 20 mg/mL in pyrogen free water (pH 6.5) using 2 ml 0.5M NaIO_4_, incubated (20°C, 60 min) and the reaction stopped with 250 μL ethylene glycol. The solution was desalted (6000 MW exclusion column equilibrated with 0.2 M borate buffer, pH 9), 1 mL was collected, and carbohydrate concentration measured per carbohydrate concentration kit instructions (Thermo Fisher Scientific, Waltham, MA). BDG conjugation fractions were pooled into three groups determined by column arrival time, representing different chain lengths for a final ratio of 1:0.5:1 (glucan, bovine serum albumin (BSA), Na cyanoborohydride). After the solution was agitated (5 days in water bath, 40°C), BDGs were separated on 6000 MW exclusion column equilibrated with 20 mM PBS pH 7.4. The heaviest carbohydrate (phenol-sulfuric acid) and protein (OD280) fractions were pooled and concentrated with a centriplus concentrator (Thermo Fisher Scientific), re-separated, and combined with the heaviest BSA. The final injection mixture contained equal volumes of three conjugate mixes. Branched glucans (scleroglucan, laminarin) were treated identically, except 2 mL of CF_3_CO_2_H (0.2M) was used for hydrolyzation.

### β-D-glucan murine exposure

On days 0 and 7, mice received intraperitoneal (i.p.) injections of glucan-bovine serum albumin conjugates (glucan-BSA) with an adjuvant of glucan (25 μg/ml) emulsified with 1 mg/mL aluminum hydroxide (alum) suspended in saline. Negative control mice were exposed to saline. On days 14-16, 21-23, and 28-30, mice previously exposed to BDGs were challenged intranasally with 25 μg BDG/mouse, suspended in 50 μL saline, while control mice received saline solution, **Appendix Fig. 1**. A sub-study was performed using BDGs autoclaved for 1h prior to nasal instillation. Procedures were performed under anesthesia, the dose of which was determined by the loss of reaction to a pinch. Euthanasia (i.p. pentobarbital, 150 mg/kg) and exsanguination were performed on day 31.

### Bronchoalveolar lavage fluid collection and analysis

Lungs were lavaged with sterile, pyrogen-free saline at a pressure of 25 cmH_2_O in 1-mL increments for a total volume of 4 mL. BAL was stored at 4°C and processed as soon as possible after collection. The BAL was centrifuged for 5 min at 800 × g, and the resulting supernatant was decanted, divided into equal volume aliquots. BAL was analyzed by multiplex immunoassay bead array systems (BioPlex, Bio-Rad, Hercules, CA) per manufacturer instructions. Antibodies were purchased from DNA Brands, Inc. (Palo Alto, CA), and biotinylated anti-TNF-α Ab from PeproTech (Thermo Fisher Scientific). BAL was analyzed for IL-6, IL-9, IL-10, IL-12p40, IL-17, CXCL-1, MCP-1/CCL-2, MIP-1α/CCL-3, MIP-1β/CCL-4 and Eotaxin.

Lung cells were analyzed for the same cytokines except MCP-1/CCL-2, MIP-1β/CCL-4 and Eotaxin. The limit of detection was 10 pg/ml for each assay. Sandwich enzyme-linked immunoassays (ELISAs) were performed using paired antibodies (eBioscience, Invitrogen, Waltham, MA) for IFN-γ, TGF-β and TNF-α. The limit of detection was 10 pg/ml for each assay. Immune cell infiltrate in BAL was assessed using previously described cell counting methods (33).

### Bronchoalveolar lavage fluid protein concentration and cell-type

The BAL cell pellet was re-suspended in Hank’s Balanced Salt Solution media with phenol red and prepared for total and differential cell counts. Total counts were performed using an improved Neubauer hemocytometer (Reichert, Buffalo, NY, USA), and the Diff Quick Stain Set (Thermo Scientific, USA) was used for staining for differential enumeration by microscopy. Total protein concentration was determined using the QuantiPro BCA assay kit (Sigma-Aldrich) by first interpolating the calibration curve using BSA standard solution (0.5 – 30 μg/ml) and concentration determined per manufacturer instructions. Briefly, 150 μL was added to the plate and incubated for 16h and absorbance was measured at 562 nm (SpectraMax Plus 384, Molecular Devices, Inc.). BAL was also analyzed for water-soluble proteins.

### Lung homogenate cell culture

Lobes from the left lung were dispersed (repeated suction/expulsion cycles through 1 ml syringe) and transferred to RPMI 1640 (GIBCO, Grand Island, NY), then passed through a 100 μm nylon cell strainer (BD Labware, Franklin, NJ). Red blood cells were lysed by hypotonic shock. Remaining cells were washed twice and re-suspended in RPMI 1640 containing 10% FCS, 25 mM HEPES buffer, 2 mM L-glutamine, 5 x 10^-5^ M β-mercaptoethanol, 1 mM sodium pyruvate, 100 U/ml penicillin, and 100 mg/ml streptomycin (GIBCO). Cell viability was quantified using trypan blue stain and determined to be >90%. Cells were cultured in triplicate (96-well microtiter plates, Corning, Cambridge, MA) (2 x 1^6^ cells/well) at 37°C and 5% CO_2_. Conditions included baseline (vehicle-treated), BDG (10 μg/ml glucan) and LPS (8 μg/ml derived from *Escherichia coli* 0111:B4, Sigma-Aldrich), or with anti-CD3 mAb (clone #2C11, ATCC) (1 μg/ml), which stimulates and activates T-cells *in vitro*. After 48h, the plate was centrifuged (800 x g, 10 min) and supernatant collected and stored at -80°C for cytokines analyses. Cytokines and chemokines evaluated included IL-4, IL-6, IL-10, IL-12/23p40, IL-17 and MIP-1α by multiplex immunoassay (Bio-Rad, Hercules, CA).

### Histopathology

Lungs of mice from each exposure group were fixed with 10% formaldehyde, embedded in paraffin, sectioned (4 μm thick), and stained with hematoxylin and eosin or Masson’s Trichrome staining. The slides were evaluated for visual abnormalities and scored for inflammation and fibrosis.

### Serum

Animals were anesthetized, euthanized, and exsanguinated by cardiac puncture to obtain whole blood. Blood from all animals of an exposure group was pooled and centrifuged (600 x g) and serum isolated and assessed for immunoglobulin E (IgE) and IgG_2a_ using sandwich ELISA (Invitrogen, Waltham, MA) according to manufacturer instructions.

### Statistics

Statistical analyses were carried out using GraphPad Prism, version 10.0.0. Values below the limit of detection were divided by √2 and cell-count values of 0 were changed to 0.5. Normality was assessed by Kolmogorov-Smirnov test. Within a given experiment, if data were normally distributed, one-way ANOVAs were performed with Šídák’s multiple comparisons test comparing control to each condition. If data were not normally distributed, a Kruskal-Wallis test was performed with Dunn’s multiple comparisons test. The number of BAL immune cells per mouse was compared to the number of cells per mL by linear regression. Sentinels, saline-exposed and saline+alum-exposed mice were compared by one-way ANOVAs or Kruskal-Wallis test with multiple comparisons. Figures denote *p* values ≤0.1 and significance was determined at *p*<0.05.

## RESULTS

### Glucan structure

Glucan linearity was assessed by nuclear magnetic resonance (NMR, **Appendix Fig. 2**). Each glucan compound used in this study has a unique branching pattern and chemical structure, **Appendix Fig. 3**. Glucan source, molecular weight, and solubility characteristics can be found in **Table 1**.

**Table 1.** Structural characteristics and solubility of glucan compounds. Molecular weight provided by BDG vendor. Linearity was assessed by NMR and solubility determined by visual assessment.

| Name | Linkage | Source | Molecular weight | Linearity | Solubility in Saline |
| --- | --- | --- | --- | --- | --- |
| Scleroglucan | (1→6)(1→3)-β-D-glucan | *Sclerotium glaucinicum* | 3,300,000 | Branched | Soluble |
| Laminarin | (1→6)(1→3)-β-D-glucan | *Laminaria digitata* | 58,500 | Linear with side branching | Soluble |
| Curdlan | (1→3)-β-D-glucan | *Alcaligenes faecalis* | 136,000 | Linear | Insoluble |
| Pustulan | (1→6)-β-D-glucan | *Umbilicaria papullosa* | 20,000 | Linear | Soluble |

### Global effects of β-D-glucan exposures

Mice were sensitized by i.p. injections to glucan conjugate emulsified with alum in saline. After two weeks, animals were challenged intranasally with individual glucans for a total of nine doses over two weeks and protein concentration and cell-type composition of BAL was analyzed. Because the concentration of immune cells per mouse was linearly correlated with the number of immune cells per mL (R2 range=0.89-1.0, *p*≤0.01, **Appendix Fig. 4**), we performed the subsequent analyses using mL as the denominator. Saline-exposed animals were used as a control unless otherwise indicated, given no significant differences were observed between saline and the saline +alum conditions, **Appendix Fig. 5**.

Pustulan exposure, but no other glucan exposures, led to the greatest BAL total protein concentration, **Fig. 1A**. Analysis of the composition of BAL cells after exposure to each glucan revealed that pustulan induced the greatest influx of inflammatory cells, significantly increasing macrophage, neutrophil, lymphocyte and total cell concentrations compared to controls (saline exposure), **Fig. 1B-E**. Scleroglucan significantly increased macrophage and total cell concentrations while curdlan increased total cell concentration in BAL compared to controls, **Fig. 1B,E**. Laminarin exposure was no different than controls for any of these outcomes.

We then tested whether glucan structure differentially affected antibody production by assessing serum immunoglobulin (IgE and IgG_2a_) concentrations after exposure to individual glucans. Both IgE and IgG_2a_ concentrations were elevated after exposure to scleroglucan and pustulan, while laminarin increased IgG_2a_ concentrations compared to saline exposures, **Figure 2A-B**. Curdlan elevated both IgE and IgG_2a_, however, responses were variable and did not reach significance.

**Figure 1.**
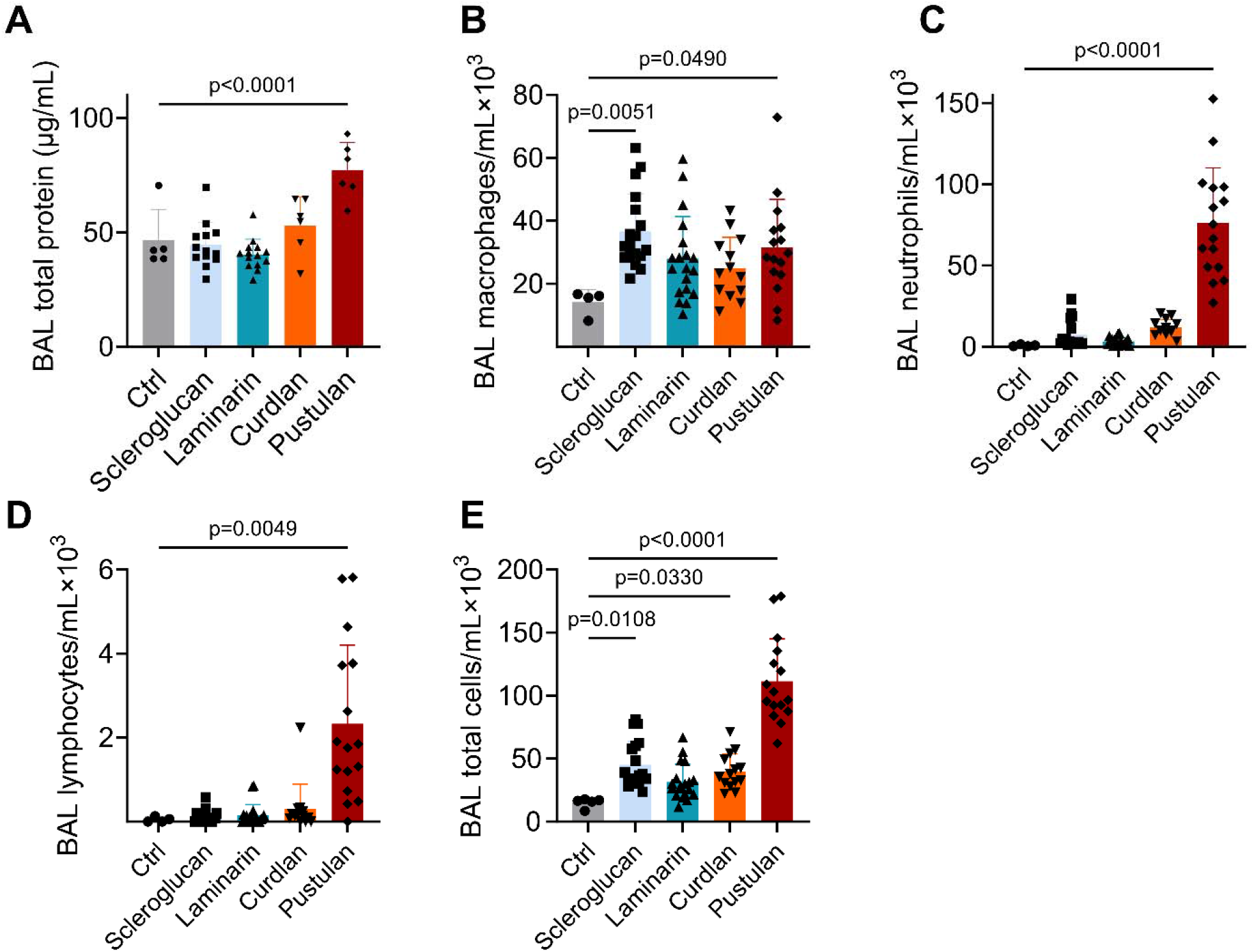
Total protein concentration and inflammatory cell infiltrate in bronchoalveolar lavage by glucan exposure. **A**. BAL protein concentration after exposure to each BDG; **B-E**. Cell concentrations in BAL after exposure to the indicated BDG. **B**. Macrophages; **C**. Neutrophils; **D**. Lymphocytes; **E**. Total cells; significance determined by comparing control to each condition by one-way ANOVA with Šídák’s multiple comparisons test (total protein, macrophages, PMNs) or by Kruskal-Wallis test with Dunn’s multiple comparisons test (lymphocytes, total cells); *n*=4-19.

**Figure 2.**
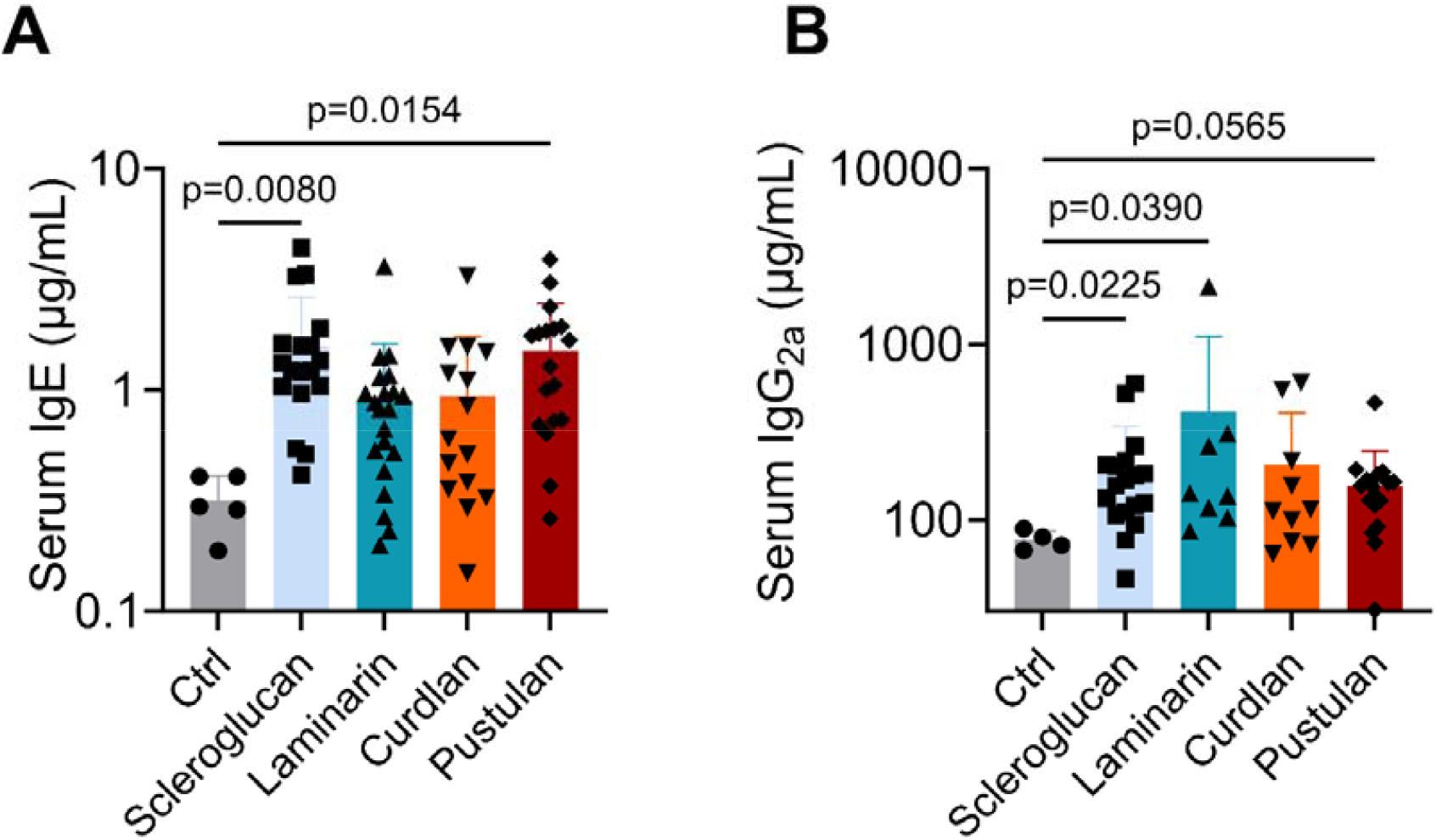
Effect of BDG exposure on serum IgE and IgG_2a_ concentrations. **A**. IgE concentration after exposure to each BDG. **B**. IgG_2a_ concentration after exposure to each BDG; significance determined by one-way ANOVA with Šídák’s multiple comparisons test (IgE) or by Kruskal-Wallis test with Dunn’s multiple comparisons test (IgG_2a_); *n*=4-21.

We analyzed BAL for the presence of Th1, Th2 and Th17 lineage cytokines to test whether exposure to unique glucan structures differentially affects cytokine production in the lungs. Pustulan induced the greatest number of changes in cytokine concentrations. Compared to saline control, pustulan increased IL-12p40, IL-17, CXCL-1, CCL-2, CCL-3, CCL-4 and decreased IL-9 and IL-10 production. Laminarin also decreased IL-9 production, while curdlan decreased IL-10 production, **Fig. 3**. IL-9 and IL-10 both have anti-inflammatory properties (34–38). No changes to IFN-γ, TGF-β and TNF-α (tested by ELISA), were observed, **Appendix Fig. 6**.

**Figure 3.**
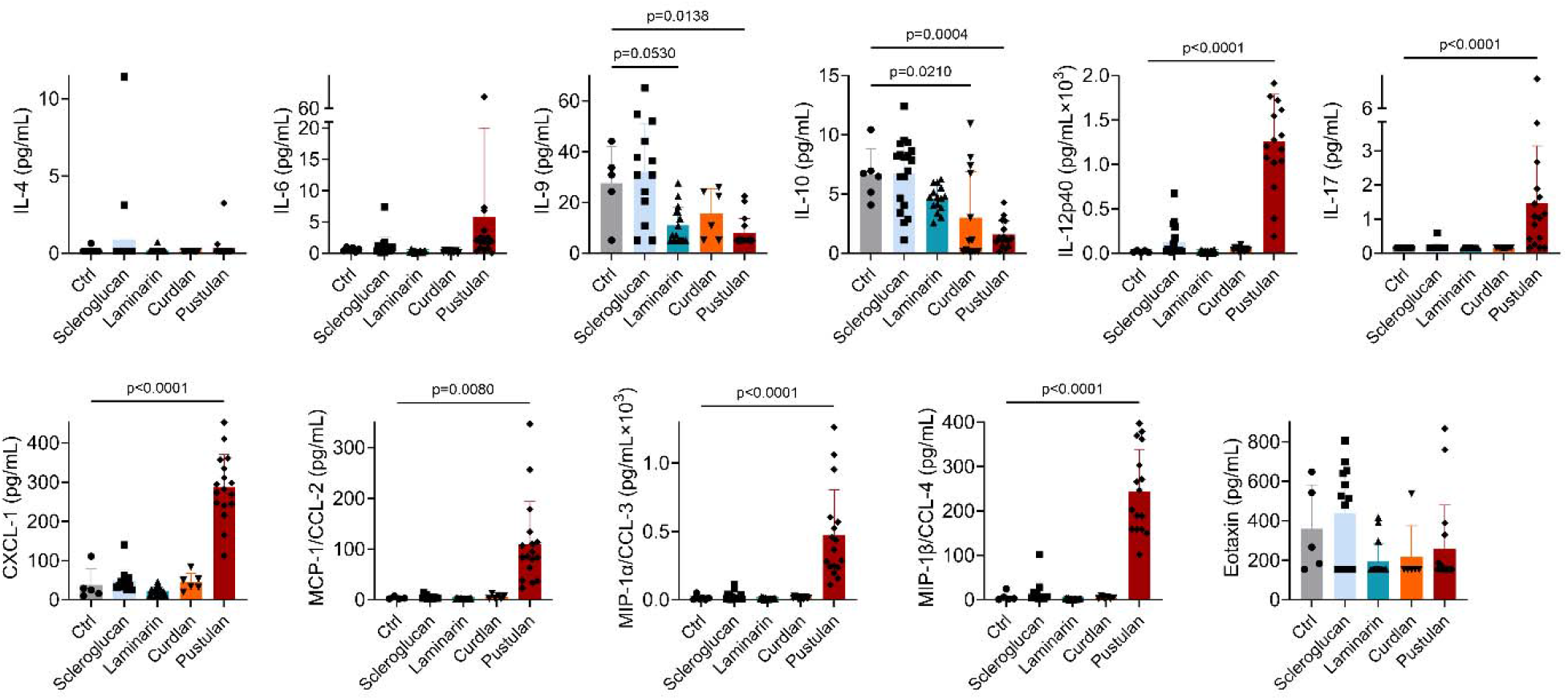
Cytokine concentration in bronchoalveolar lavage by BDG exposure. IL-4, IL-6, IL-9, IL-10, IL-12p40, IL-17, CXCL-1, MCP-1/CCL-2, MIP-1α/CCL-3, MIP-1β/CCL-4 and Eotaxin concentration in BAL; significance determined by comparing control to each condition by one-way ANOVA with Šídák’s multiple comparisons test or by Kruskal-Wallis test with Dunn’s multiple comparisons test depending on Kolmogorov-Smirnov test results; *n*=5-19.

### Effect of β-D-glucan solubility on murine lungs

Indoor BDG exposure quantification often relies on first heating dust samples prior to analysis (39–41), which increases their solubility. Soluble BDG is a Dectin-1 antagonist, while particulate BDG is a Dectin-1 agonist. Soluble BDGs might, however, agonize other non-dectin-1 β-glucan PRRs (14, 15, 42). Therefore, we sought to analyze the effect of solubility on lung inflammation. We performed a sub-study on the effect of each compound after it was autoclaved (1h) to assess the pro-inflammatory effect. Curdlan and pustulan heating markedly increased recruitment of inflammatory cells to the lung, **Fig. 4A-B, D**. Contrary to prior findings, heated scleroglucan no longer increased macrophage or total cell concentrations in BAL, **Fig. 4A, D**. Although lymphocyte response was quite variable, heating of pustulan decreased the migration of lymphocytes into BAL by approximately one order of magnitude, **Fig. 4C**.

**Figure 4.**
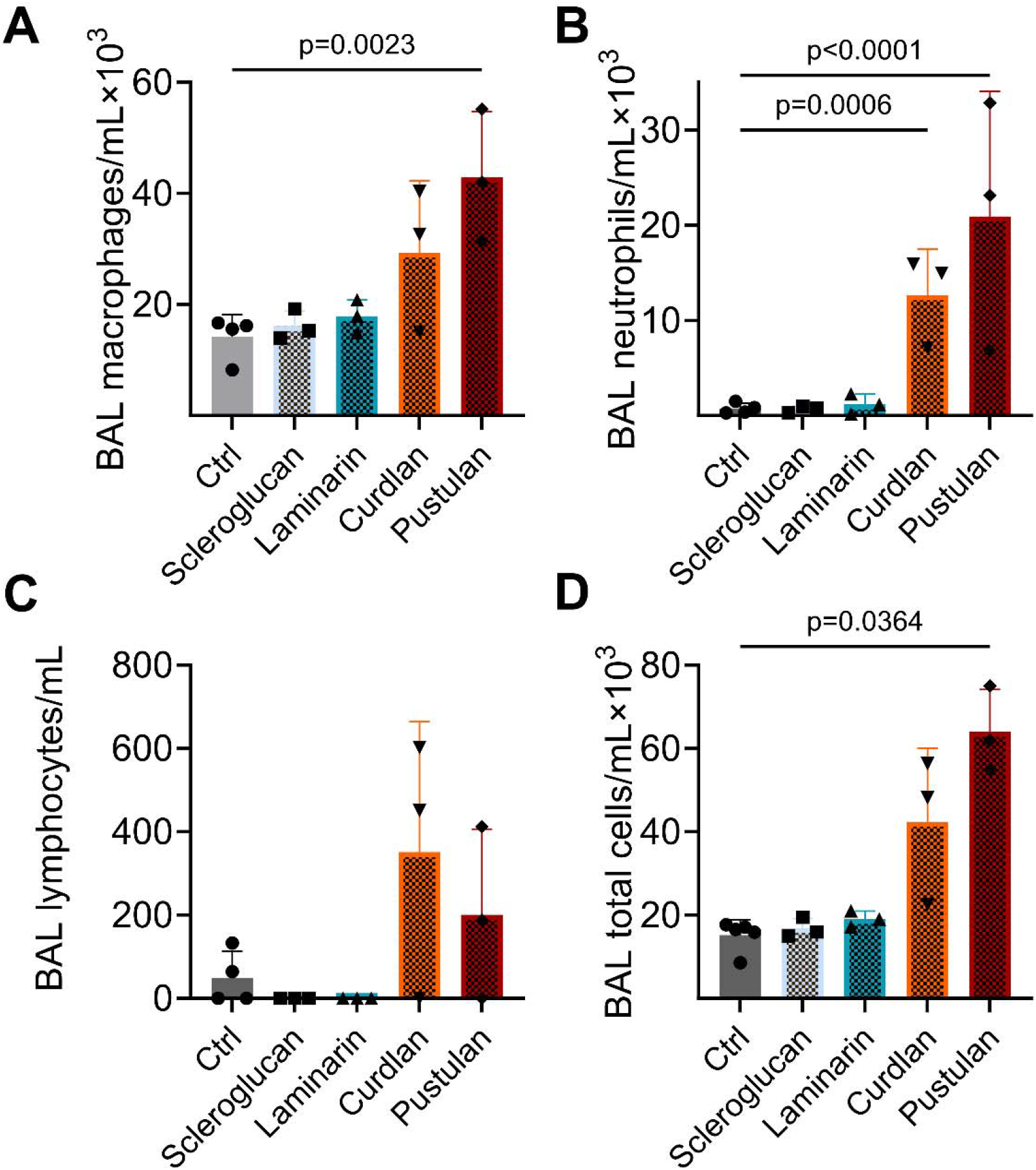
Effect of glucan heating on inflammatory cell infiltrates in bronchoalveolar lavage by BDG exposure. **A-D**. Cell concentrations in BAL after exposure to each BDG. A. Macrophages; **B**. Neutrophils; **C**. Lymphocytes; **D**. Total cells; significance determined by comparing control to each condition by one-way ANOVA with Šídák’s multiple comparisons test (macrophages, PMNs) or by Kruskal-Wallis test with Dunn’s multiple comparisons test (lymphocytes, total cells); *n*=3-5.

Next, we assessed the effect of glucan solubility on BAL cytokine expression. While cytokines were typically below the limit of detection, IL-10, IL-12p40 and CCL-3 concentrations were detectable in BAL, **Fig. 5**. Heated pustulan increased CCL-3 concentration significantly compared to control, **Fig. 5C**. Heating of glucans had no effect on TNF-α (by ELISA), **Appendix Fig. 6B**.

**Figure 5.**
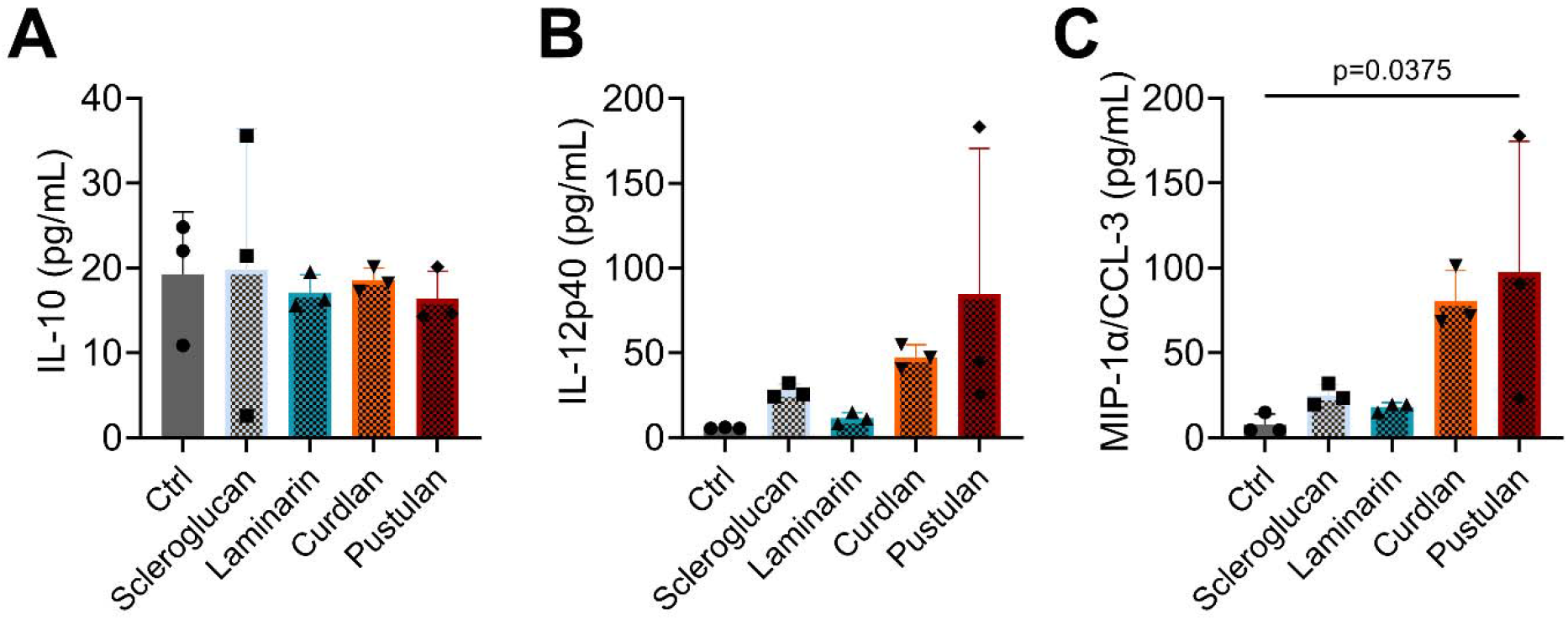
Effect of heating on cytokine concentration in bronchoalveolar lavage by glucan exposure. **A**. IL-10; **B**. IL-12p40; **C**. MIP-1α/CCL-3 concentration in BAL; significance determined by comparing sentinel controls to each condition by one-way ANOVA with Dunnett’s multiple comparisons test; *n*=3 per condition.

We further tested the effect of glucan solubility on BAL cytokine concentrations by comparing unheated and heated glucans for IL-10, IL-12p40 and MIP-1α/CCL-3, **Fig. 6**. Typically, heating of BDGs increased the concentrations of cytokines present in BAL. Heating of each glucan significantly increased the concentration of IL-10, **Fig. 6A**, while heated laminarin and pustulan decreased IL-12p40, **Fig. 6B**. We observed differential effects of BDG heating on MIP-1α/CCL-3. Heated laminarin and curdlan increased MIP-1α/CCL-3, while pustulan decreased MIP-1α/CCL-3 (trend), **Fig. 6C**.

**Figure 6.**
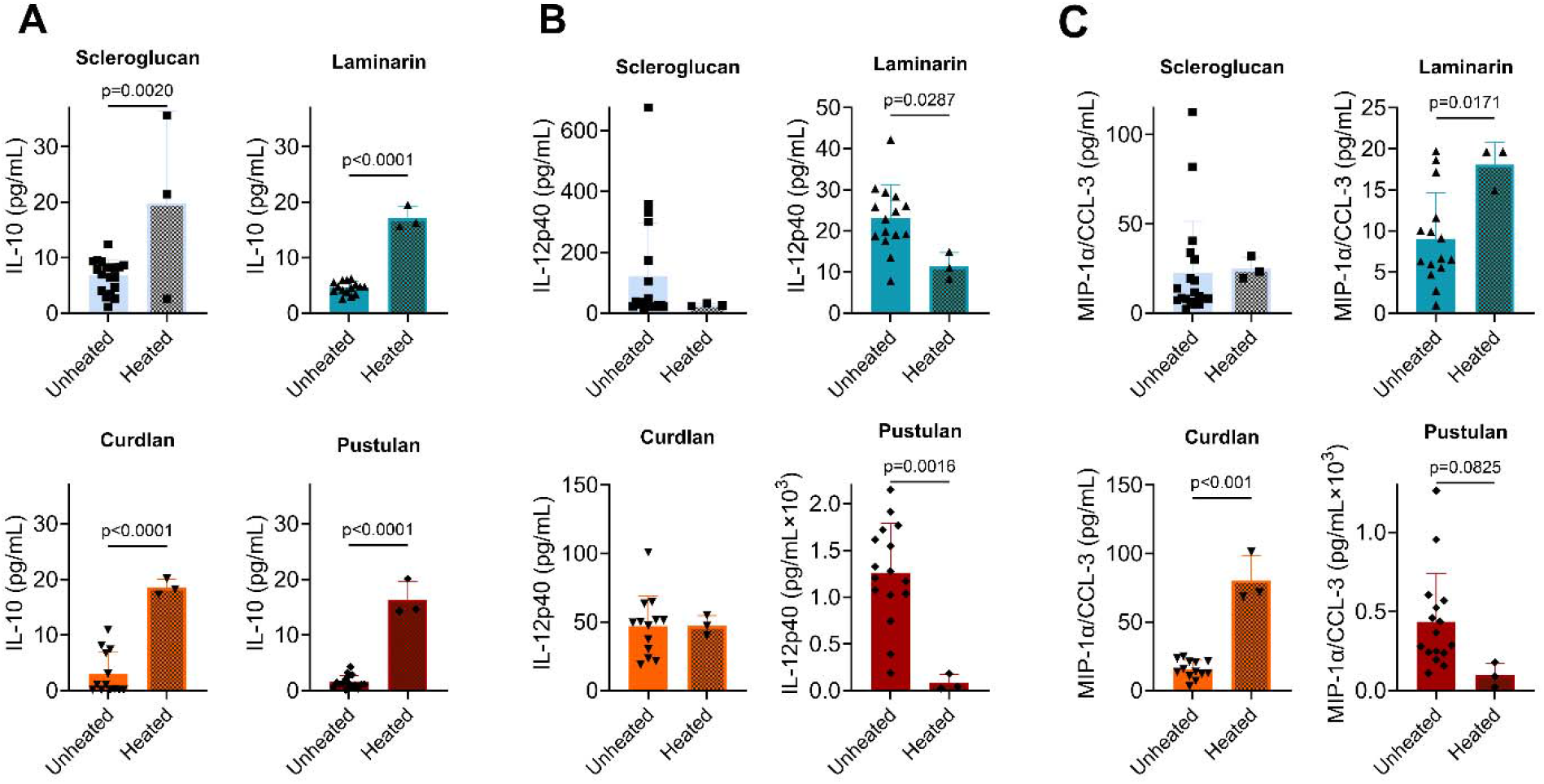
Effect of glucan heating on cytokine concentration in bronchoalveolar lavage by BDG exposure. **A**. IL-10 concentration in BAL. **B**. IL-12p40 concentration in BAL. **C**. MIP-1α/CCL-3 concentration in BAL; significance determined by unpaired *t*-test; *n*=3-19.

Next, we analyzed lung histology in animals exposed to BDG with or without heating. Overall, all glucans induced neutrophil and lymphocyte migration to the airspace. Within the scleroglucan-exposed group, macrophages were noticeably larger, while pustulan induced the greatest infiltrate. Heating of insoluble, linear curdlan increased inflammatory infiltrate while unheated glucans otherwise had the greatest effect, **Fig. 7**.

**Figure 7.**
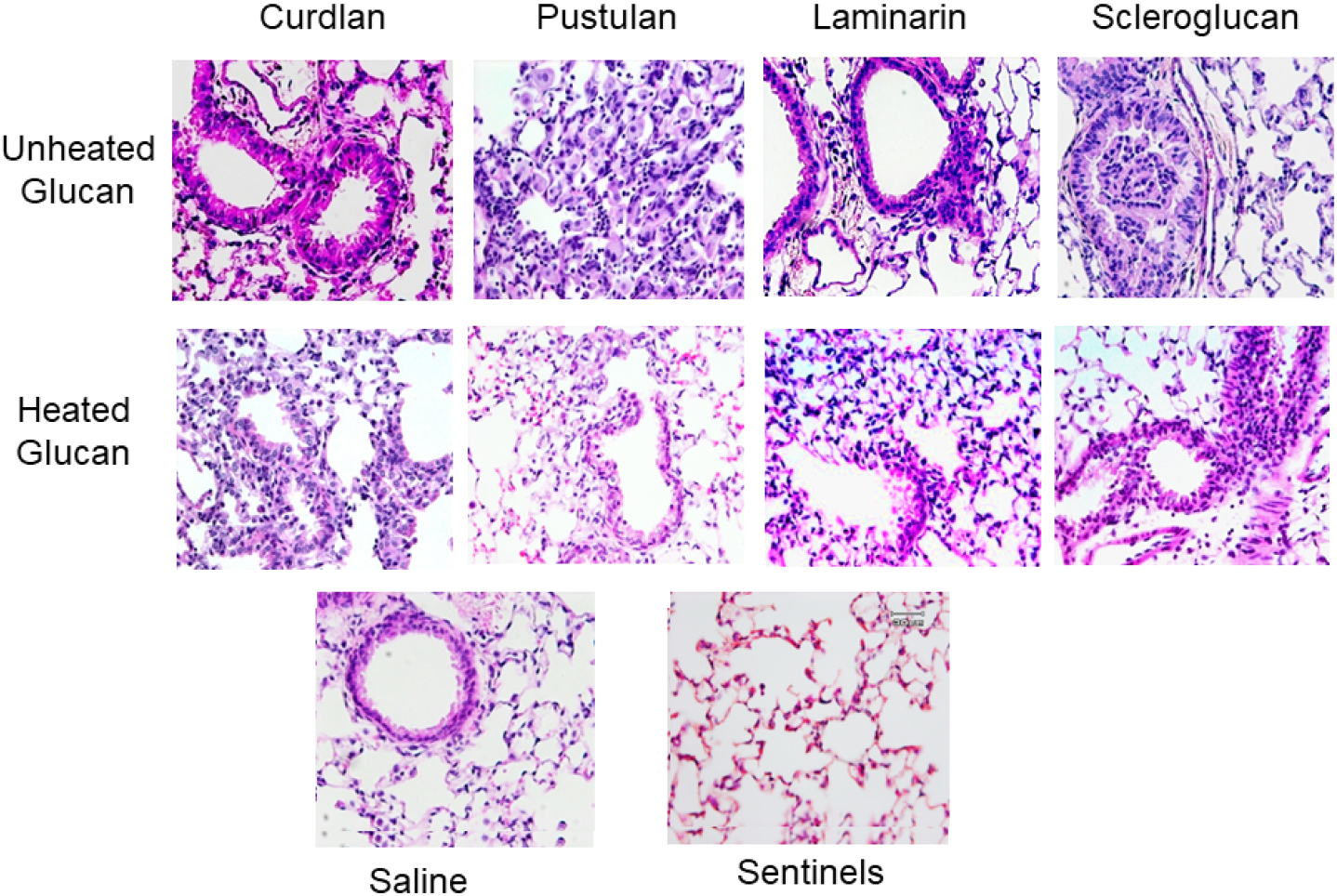
Histopathology of BDG-exposed lungs. Mice were exposed to unheated or heated glucan compounds and lungs were stained with hematoxylin and eosin stain; representative photomicrographs selected from separate experiments representing similar results per condition. All images presented with identical brightness, contrast, and magnification (40x).

To assess whether BDG exposure (unheated) led to fibrosis, we stained lung sections with Masson’s trichrome. We observed minor evidence of fibrosis in the scleroglucan group, and significant fibrosis in the pustulan group, **Fig. 8B, D**.

**Figure 8.**
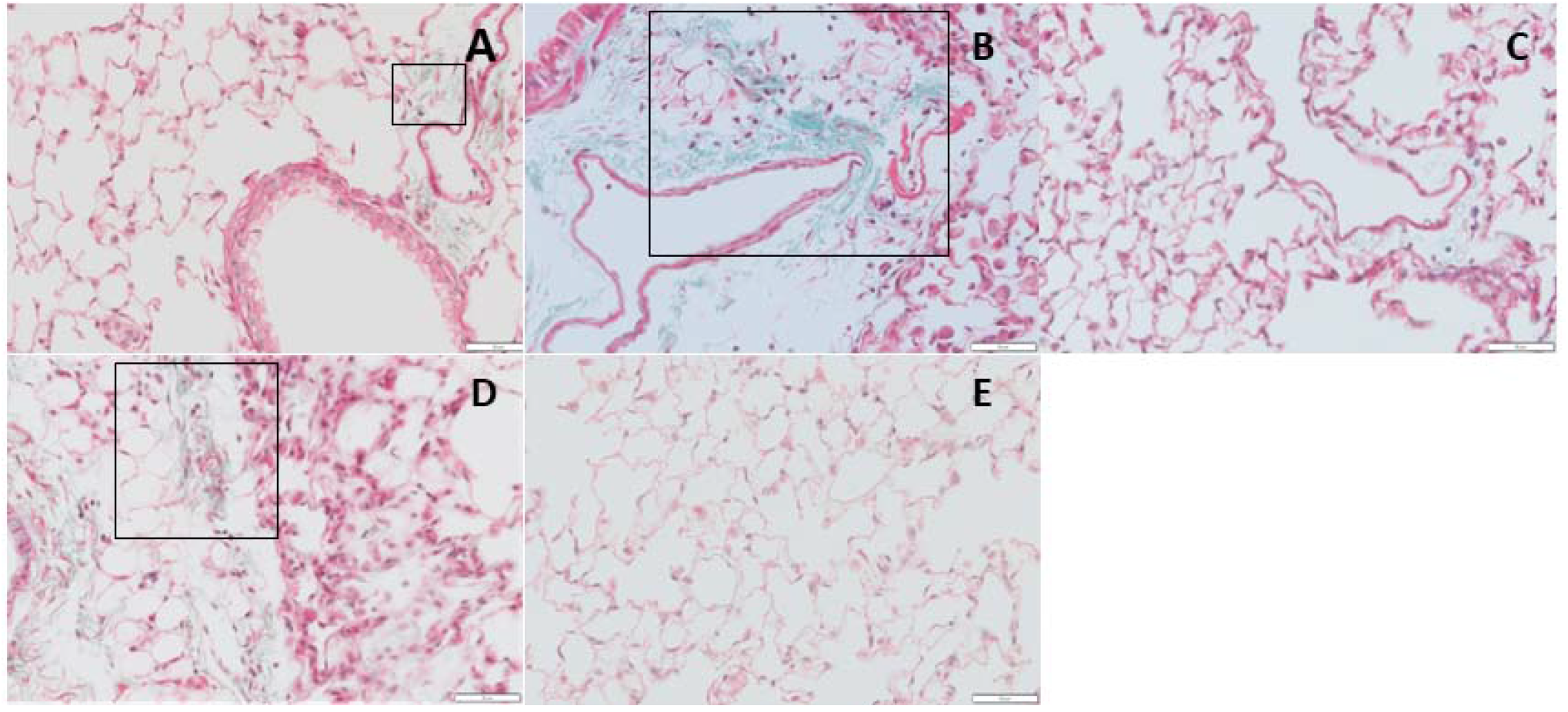
Micrographs of lung sections stained with Masson’s trichrome after mice were exposed to unheated glucan. **A**. Curdlan exposure led to very little fibrosis (+/-); **B**. Pustulan exposure led to significant fibrosis (+/+); **C**. Laminarin exposure led to very little fibrosis (+/-), not shown; **D**. Scleroglucan exposure led to minor fibrosis (+); **E**. Saline-exposed mice did not have fibrosis (-). Areas of significant lung fibrosis are identified by black box; magnification=40x.

### *Ex vivo* lung homogenate cell stimulation

We next tested whether primary sensitization to BDGs affected *ex-vivo* lung cell cytokine production upon secondary stimuli. Lobes from the left lung were dissected, homogenized, and cultured in triplicate, then exposed to vehicle, BDG, LPS, or CD3 mAb (to stimulate T-cells); the cell supernatant was collected after 48h, and cytokines assessed.

Lipopolysaccharide (LPS) exposure to cultured airway cells previously sensitized (*in vivo*) increased IL-4 (1° stimuli: curdlan, pustulan), IL-6 (all), IL-10 (pustulan), IL-12p40 (curdlan), and MIP-1α/CCL-3 (pustulan) cytokine concentrations, **Fig. 9A-D, F**. We next assessed T-cell specific cytokine induction by addition of a CD3 antibody, observing differential effects based on initial BDG stimulation. For example, compared to baseline, IL-4 increased across all conditions, however not significantly; IL-6 increased (1° stimulus: pustulan), IL-12p40 decreased (pustulan), IL-17 increased (all trend) and MIP-1α/CCL-3 increased (pustulan trend), **Fig. 9**. Secondary exposure to BDGs did not increase cytokine expression, except in the case of curdlan, which increased IL-4, IL-6, IL-10, IL12p40 and MIP-1α/CCL-3 production compared to control, **Appendix Fig. 7**.

**Figure 9.**
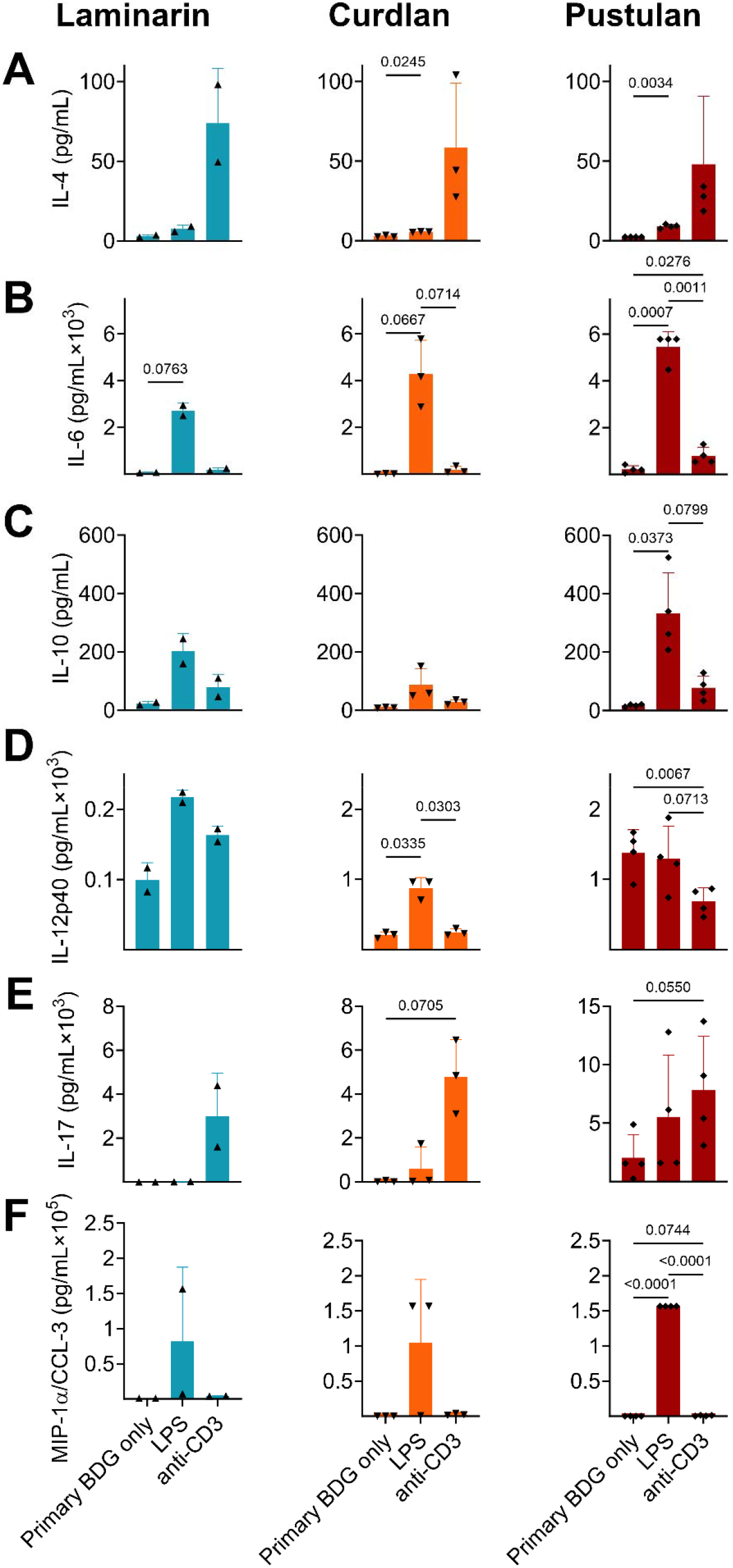
Effect of secondary stimuli on cultured lung homogenate cells. Effect of LPS and anti-CD3 antibody on **A**. IL-4, **B**. IL-6, **C**. IL-10 D. IL-12p40, **E**. IL-17, **F**. MIP-1α/CCL-3 concentrations in cell supernatant; significance determined by repeated measures one-way ANOVA with Tukey’s multiple comparisons test; *n*=2-4, performed in triplicate.

## DISCUSSION

We tested the effect of four structurally distinct β-D-glucans on murine immune response, finding that soluble (1→6)-linear BDG pustulan exposure led to the most pronounced pro-inflammatory effects. Pustulan is from lichen; exposure to which likely occurs in occupations working with wood dust and in the lumber/forestry industry, which increase the risk of pulmonary fibrosis and sarcoidosis (43, 44). Farmers are potentially exposed to scleroglucan when harvesting soybeans (45). Curdlan originates from environmental bacteria, making exposure to it common. Laminarin originates from algae making exposure potentially less likely.

Pustulan increased total BAL protein concentration, immune cell infiltrate, cytokine concentrations, IgE and IgG_2a_, and eventual fibrosis observed by histology, compared to control **Figs. 1-5, 8**. The neutrophil and lymphocyte migration into the airspace observed in lung micrographs were reminiscent of a murine asthma model (46), however, our cytokine data supports a mixed Th1/17 immune response to BDGs, **Fig. 3**. Distinct mold-induced phenotypes have been reported; for example, sarcoidosis which is typically considered a Th1 and/or Th17-dominate disease (47, 48), has been identified as elevated in workers exposed to a water-damaged building but this same building also led to increased asthma incidence and symptoms, likely indicative of gene-environment interactions (4, 49).

Allergic asthma is an inflammatory lung disease characterized by eosinophil infiltrate and bronchial hyperreactivity (50, 51). Scleroglucan (1→ 3, 1→ 6)-b-D-glucan exposure in asthmatic children has been shown to decrease FEV_1_ and greatly increases the odds of emergency visits for asthma (5). In this work, scleroglucan induced certain hallmarks of asthma, including increased macrophages (number and size) in the airspace and minor fibrosis, **Fig. 1B, E**; **Figs. 7-8** and increased serum IgE, **Fig. 2A** (52). However, Th1 IgG_2a_ also increased **Fig. 2B**.

While allergic disease is predominated by T cell differentiation into Th2 cell response, Th1 cytokine, IFN-γ, can also potentiate lung injury induced by Th2 cytokine IL-13 (53). Interplay between Th1 and Th2 cytokines can result in allergic disease pathology. Surprisingly, pustulan also reduced Th2 cytokine IL-9 (54) and suppressor IL-10 concentrations in BAL, **Fig. 3**, suggesting a change from Th2 towards Th1 or Th17. While we did not observe significant differences in IFN-γ or TNF-α production after exposures, **Appendix Fig. 6**, curdlan and pustulan tended to increase IFN-γ concentrations (both *p*<0.19). However, a lack of IFN-γ response is consistent with findings from Hadebe *et al*., who found IFN-γ did not respond to a co-BDG + house dust mite allergen model, (55).

We performed a sub-study to evaluate the effect of glucan solubility on lung inflammation given BDGs exist in different structures, which may alter a host’s physiological response. Glucan solubility often increased cytokine concentrations in BAL, **Fig. 6**. Further, soluble (at baseline) pustulan and scleroglucan induced lung fibrosis, which was exaggerated in the pustulan-exposed group, **Fig. 8**. Pustulan also significantly increased IL-17 concentrations in the airspace, **Fig. 3**, which is thought to play a direct role in lung fibrosis (56, 57). We believe the interaction between IL-17 and TLR4 contribute to the fibrotic phenotype we observed for the following reasons. Dectin-1 plays a protective role in fibrosis by suppressing TLR4 activation (58). Dectin-1 signaling is effectively halted when soluble BDGs are detected by innate immune cells (14, 58). The absence of Dectin-1 signaling upon exposure to soluble glucans likely increases the role of other BDG recognition components of the plasma membrane, such as TLR4 and lactosylceramide (15, 59). Further, the presence of IL-17, observed in BAL of the pustulan-exposure group, **Fig. 3**, increases TLR4 expression (60). TLR4 activation is required for IL-17 induced effects, including neutrophil infiltrate (60). Abrogation of Dectin-1 signaling likely in part drives the increased fibrosis we observed after exposure to soluble glucans.

Pustulan and scleroglucan also increased systemic inflammatory markers (IgE and IgG_2a_), **Fig. 2**. Increased serum IgG has been observed in a small cohort of patients with pulmonary fibrosis (61). Bronchoalveolar lavage from mice exposed to both soluble glucans had unique cytokine responses, perhaps indicative of distinct etiologies of lung fibrosis. Other, pro-fibrotic mediators not measured in our study may also contribute to the development of pulmonary fibrosis we observed.

Pro-fibrotic exposures (pustulan and scleroglucan) also increased BAL cell counts and macrophage concentrations. Macrophages are increasingly recognized for their role in development of pulmonary fibrosis (62–64) with excessive M2 macrophages playing a pro-fibrotic role (65). Macrophage response to BDGs has been demonstrated to be independent of the classic BDG receptors Dectin-1 and TLR2 (66, 67); because Dectin-1 is not independently responsible for Th2 sensitization in a BDG (+house dust mite) mouse model (55), macrophage recruitment and signaling might explain the fibrotic phenotype observed after exposure to these BDGs.

Pro-fibrotic exposures (pustulan and scleroglucan) also increased BAL cell counts and macrophage concentrations. Macrophages are increasingly recognized for their role in development of pulmonary fibrosis (62–64) with excessive M2 macrophages playing a pro-fibrotic role (65). Macrophage response to BDGs has been demonstrated to be independent of the classic BDG receptors Dectin-1 and TLR2 (66, 67); because Dectin-1 is not independently responsible for Th2 sensitization in a BDG (+house dust mite) mouse model (55), macrophage recruitment and signaling might explain the fibrotic phenotype observed after exposure to these BDGs.

Isolated lung cells that were exposed to secondary stimulation *in vitro* had exaggerated cytokine response to LPS, indicating the importance of BDG priming in TLR4-mediated (68) inflammatory response, **Fig. 9**. BDG exposure is known to enhance immune response and protect against bacterial infection (16), likely explaining the excessive cytokine release observed after both stimuli. In most experiments, pustulan induced the greatest inflammatory response, however, secondary curdlan exposure increased all cytokine expression in isolated lung cells *in vitro* except IL-17, **Appendix Fig. 7**. This may indicate lung cell-specific sensitization to curdlan. Further, the timepoint of BAL collection (31 days post initial exposure) may not have captured the peak inflammatory response to individual BDGs.

When T-cells were stimulated by CD3 mAb, IL-17 tended to increase *in vitro*, **Fig. 9E**. IL-17 is known to protect against fungal and bacterial infection through neutrophil recruitment, increased antimicrobial peptide production, and improved barrier protection. This is likely due to increased T-cell response, which is the primary producer of IL-17; however the effect is not limited to T-cell specific response given its can be produced by cytokines (IL-1β, IL-23), innate lymphoid cells, natural killer cells, and mast cells (69). Because IL-4 also tended to increase under these conditions, it is possible mast-cells also contribute to the increased cytokine production observed with the CD3 antibody (70). Secondary BDG stimulation of cultured lung homogenate did not enhance baseline cytokine response except in the case of curdlan, **Appendix Fig. 7**, which significantly increased all measured cytokines and chemokines except IL-17. It especially increased MIP1a/CCL-3, which hones dendritic cells to drain lymph nodes and is important in lung diseases, including sarcoidosis (71, 72).

Study strengths include the systematic use of characterized unique BDGs in an animal model of inflammation and the identification of one extremely pro-inflammatory BDG (pustulan) leading to the development of lung fibrosis. This study has several limitations. Cytokines were only measured in the lung compartment and not systemically, responses in murine lungs may not equivate to human responses, and BAL macrophage lineage was not assessed.

Overall, we observed a strong Th1 and Th17 BAL response to BDG exposure, yet serum IgE was elevated, indicative of Th2 allergen response. Histological examination demonstrated the lungs had a fibrotic appearance. BDG solubility likely increased the pro-inflammatory effects of exposures. *In vitro* secondary stimulation (LPS or BDG) of glucan-primed cells magnified the inflammatory response, especially to secondary curdlan stimulation, demonstrating the importance of prior BDG immune priming. Further, this work demonstrated structural differences in BDGs result in unique immunological responses in murine lungs. Exposures to fungi should be considered in diverse lung disease phenotypes.

## ACKNOWLEDGEMENTS

We would like to thank Dr. David L. Williams (East Tennessee State University) for his generous donation of scleroglucan. A big thanks to Andrea Adamcakova-Dodd (University of Iowa), who was generous with her time and imaging expertise. Lastly, Hui Wang provided access to the chemical structure software to map BDG structures.

