## Appendix material for "(1→3)-β-D-glucan derivatives with unique structural properties differentially affect murine lung inflammation and histology"

**MATERIALS AND METHODS**

Determination of the degree of glucan branching
The degree of branching of selected glucans was determined by using nuclear magnetic resonance (NMR). NMR experiments were performed using a Bruker FT-NMR spectrometer DMX 600 resonating at 600.13 MHz for 1H at the University of Iowa NMR facility core. Glucan compounds were dissolved in deuterium oxide (D2O) or mixtures of PBS-D2O. Two-dimensional magnitude correlation spectroscopy (COSY) experiments were performed to assign the 1H resonance.

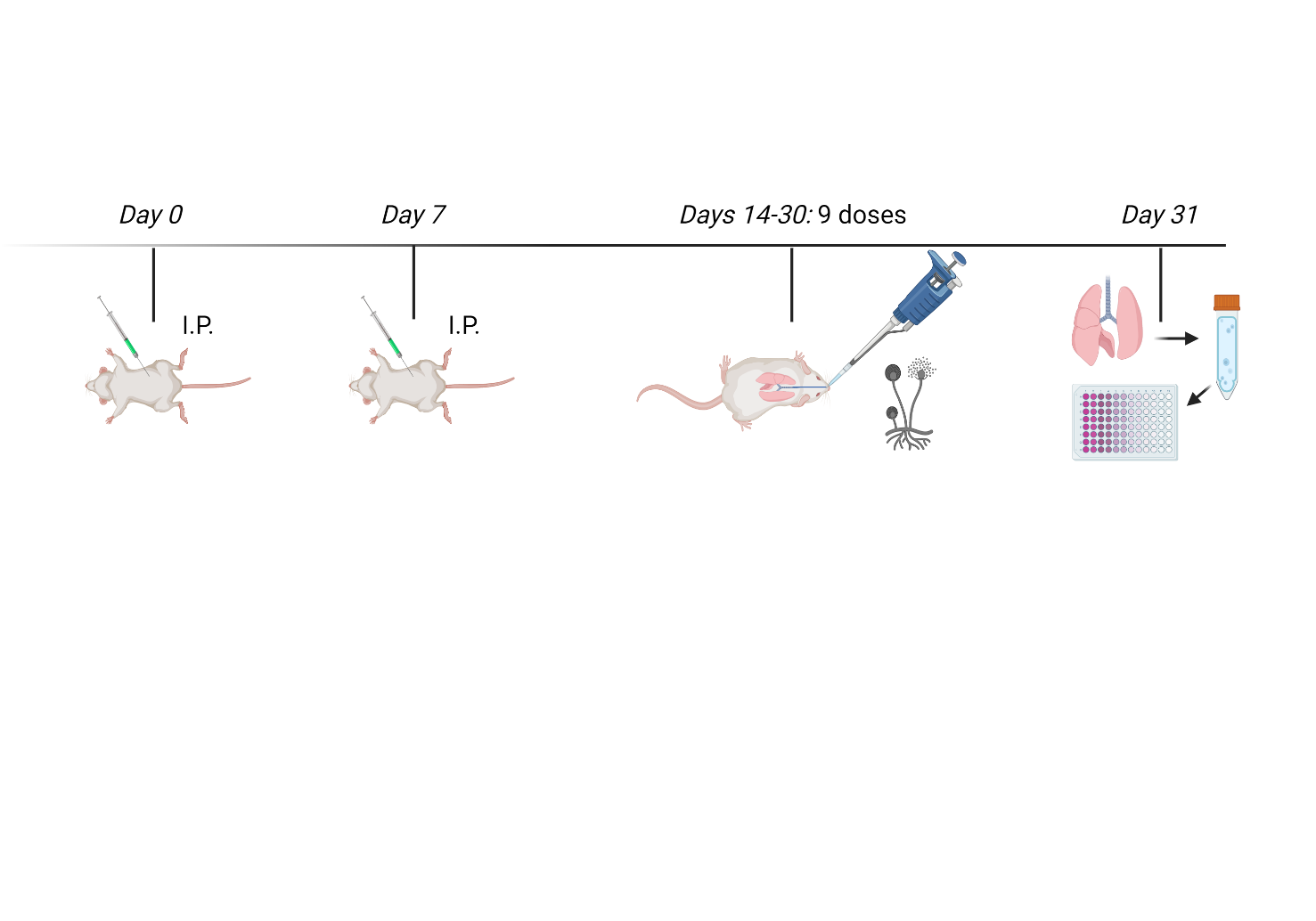

**Figure 1. Schematic of study design protocol.** Mice received intraperitoneal injections of BSA-glucan conjugates at days 0 and 7, while negative control mice were exposed to saline. On days 14-16, 21-23, and 28-30, mice were exposed to BDGs (25 µg BDG/mouse) intranasally, while control mice received saline solution. Harvest was performed on day 31; created with BioRender.com.

**RESULTS**

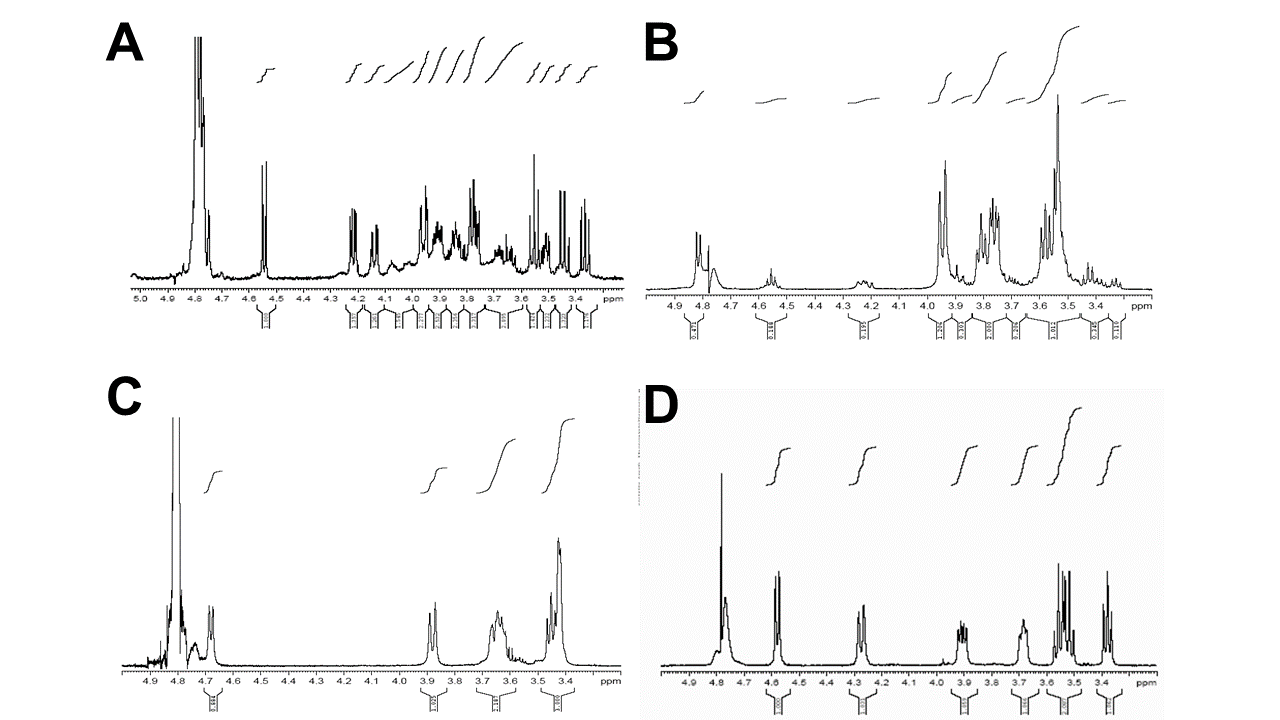

**Figure 2. Glucan compound structure assessed by nuclear magnetic resonance imaging.** Glucans were suspended in deuterium oxide and assessed by 1H NMR spectra; **A.** Scleroglucan **B.** Laminarin **C.** Curdlan, and **D.** Pustulan.

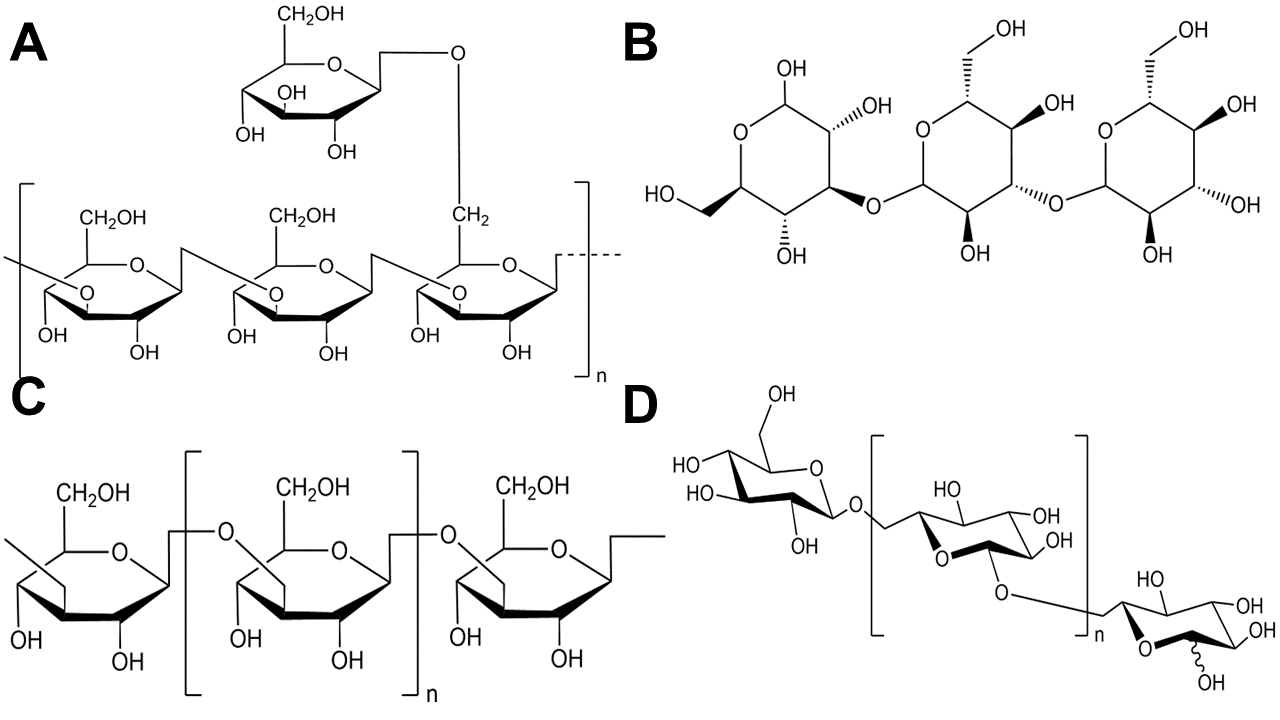
**Figure 3. A.** Scleroglucan (1→3) (1→6)-highly branched BDG. **B.** Laminarin ((1→3) (1→6)-branched BDG. **C.** Curdlan (1→3)-linear BDG. **D.** Pustulan (1→6)-linear BDG.

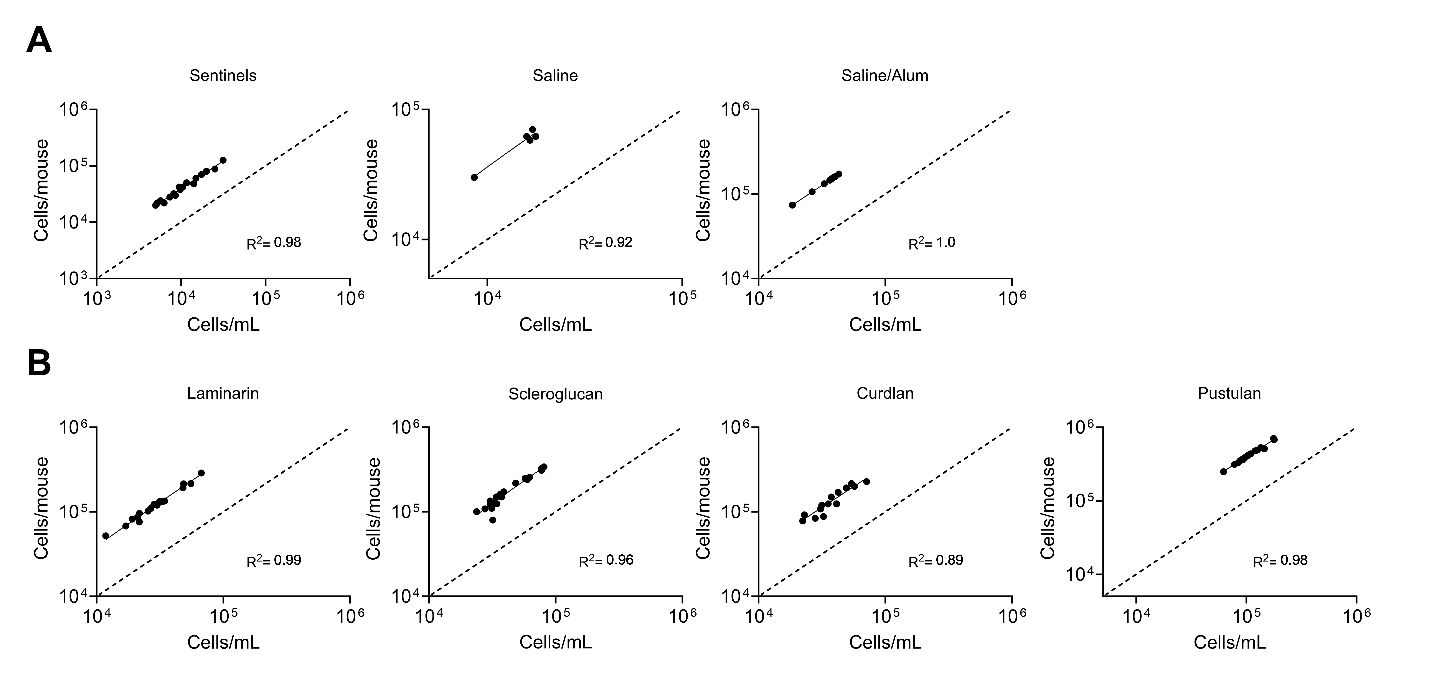

**Figure 4. Correlation between cells per mouse and cells per milliliter of bronchoalveolar lavage. A.** Comparisons between three control groups (sentinels, saline-exposed mice, and saline+alum exposed mice) demonstrate linearity between outcomes; simple linear regression R^2^>0.92; p<0.01 all conditions. **B.** Comparison of correlation between cells/mouse vs. cells/mL for each BDG exposure; simple linear regression R^2^>0.89; p<0.001 all conditions.

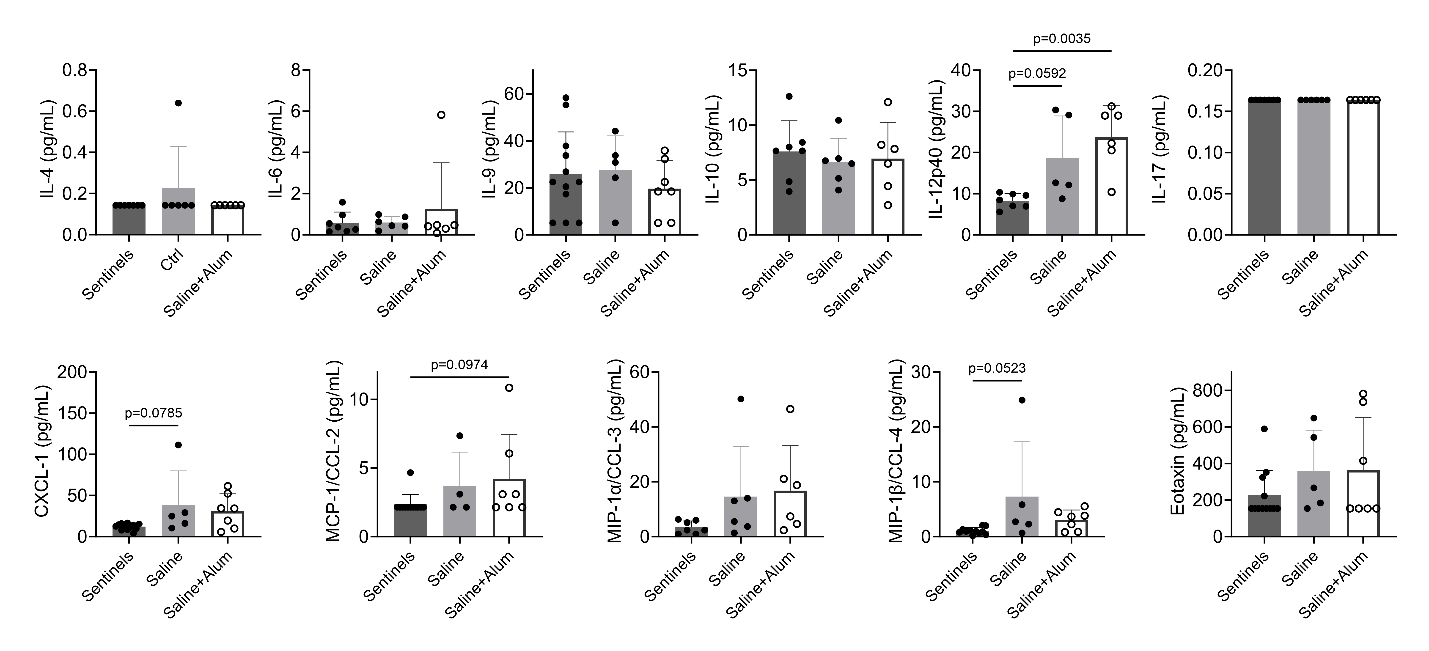

**Figure 5. Comparisons of cytokine production between control conditions.** Bronchoalveolar lavage cytokine production was similar between sentinels, saline, and saline+alum conditions, with no significant differences observed between saline or saline+alum exposed animals. Ordinary one-way ANOVA or Kruskal-Wallis tests were performed based on Kolmogorov-Smirnov normality test results, *p* values <0.1 displayed.

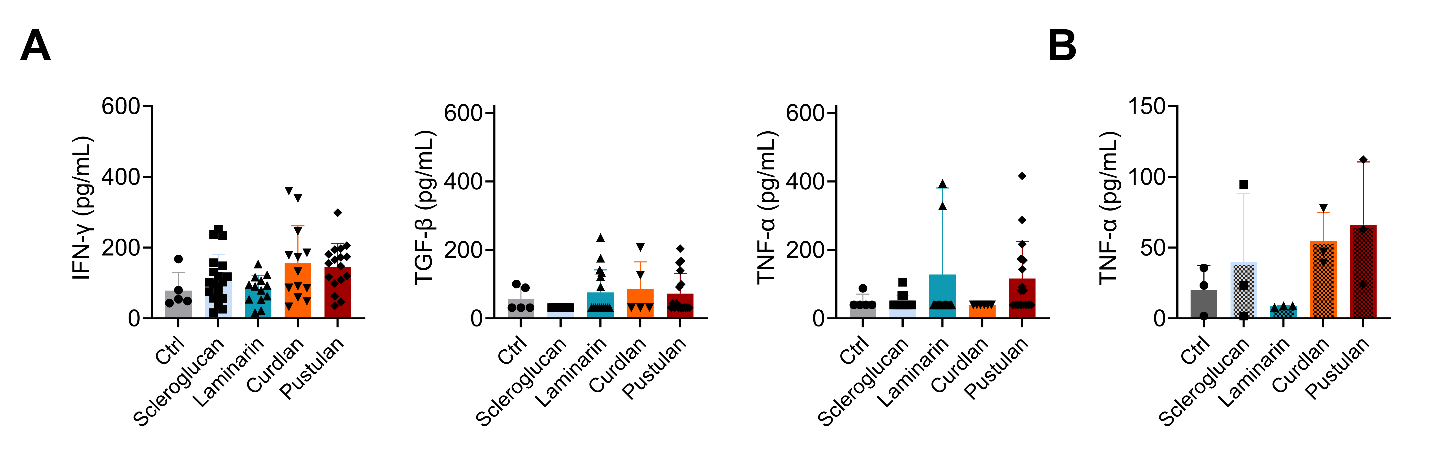
**Figure 6. Cytokines identified by ELISA. A.** No significant differences were observed between IFN-γ, TGF-β and TNF-α vs. saline controls, measured by ELISA in BAL after exposure to unheated glucans. **B.** No significant differences were observed in TNF-α vs. sentinel controls, measured by ELISA after mice were exposed to solubilized (heated) glucans in sub-study; ordinary one-way ANOVA or Kruskal-Wallis tests were performed based on Kolmogorov-Smirnov normality test results all *p*>0.1.

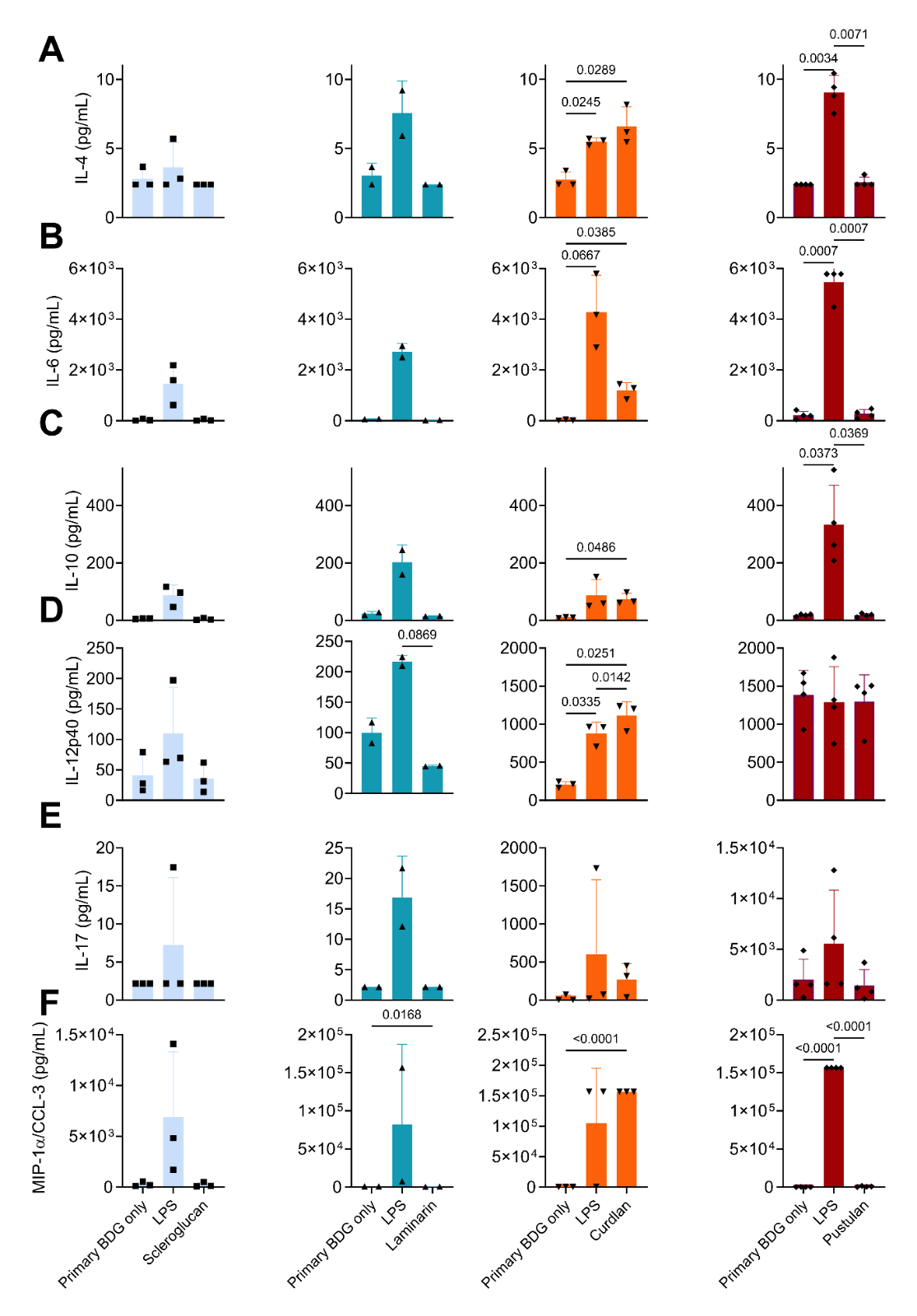

**Figure 7. Secondary curdlan stimulation in isolated lung cells results in increased cytokine expression. A.** Effect of secondary scleroglucan, laminarin, curdlan or pustulan stimulation on IL-4 production by isolated lung cells. **B.** Effect of secondary BDG stimulation on IL-6 production by isolated lung cells. **C.** Effect of secondary BDG stimulation on IL-10 production by isolated lung cells. **D.** Effect of secondary BDG stimulation on IL-12p40 production by isolated lung cells. **E.** Effect of secondary BDG stimulation on IL-17 production by isolated lung cells. **F.** Effect of secondary BDG stimulation on MIP-1α/CCL-3 by isolated lung cells; significance determined by repeated measures one-way ANOVA with Tukey’s multiple comparisons test; *p*<0.10 shown.
